# A new FKBP51-GR-p23-Hsp90_2_ multi-cochaperone complex identified and characterized by site-specific in-cell photocrosslinking

**DOI:** 10.1101/2025.06.09.658717

**Authors:** Martha C. Taubert, Angela Kühn, Asat Baischew, Yannik Käseberg, Janna Betschinske, Felix Hausch

## Abstract

Glucocorticoid receptor (GR) activity and maturation are closely regulated by Hsp90 co-chaperones such as FKBP51, FKBP52 and p23. These co-chaperones bind to and regulate the GR, but their mutual interplay, the details of the interactions between them and their temporal dynamics are still unclear.

Here we utilized UV-inducible crosslinking in living cells to map the interaction site of p23 and the GR, as well as p23 and FKBP51 at a single residue resolution. Surprisingly, we detected a novel multi-co-chaperone complex consisting of the GR, p23, FKBP51 and Hsp90, where both FKBP51 and p23 bind to the GR simultaneously. In this complex, FKBP51, but not the close homolog FKBP52, stabilizes the GR-p23 interaction. This is mediated in part by direct contacts between the FK1 domain of FKBP51 and the C-terminus of p23. Our findings refine the state of GR prior to activation, add a new layer of GR regulation by chaperones, provide evidence for the functional differences between FKBP51 and FKBP52, and underscore the power of photocrosslinking to functionally probe protein-protein contacts inside living cells.

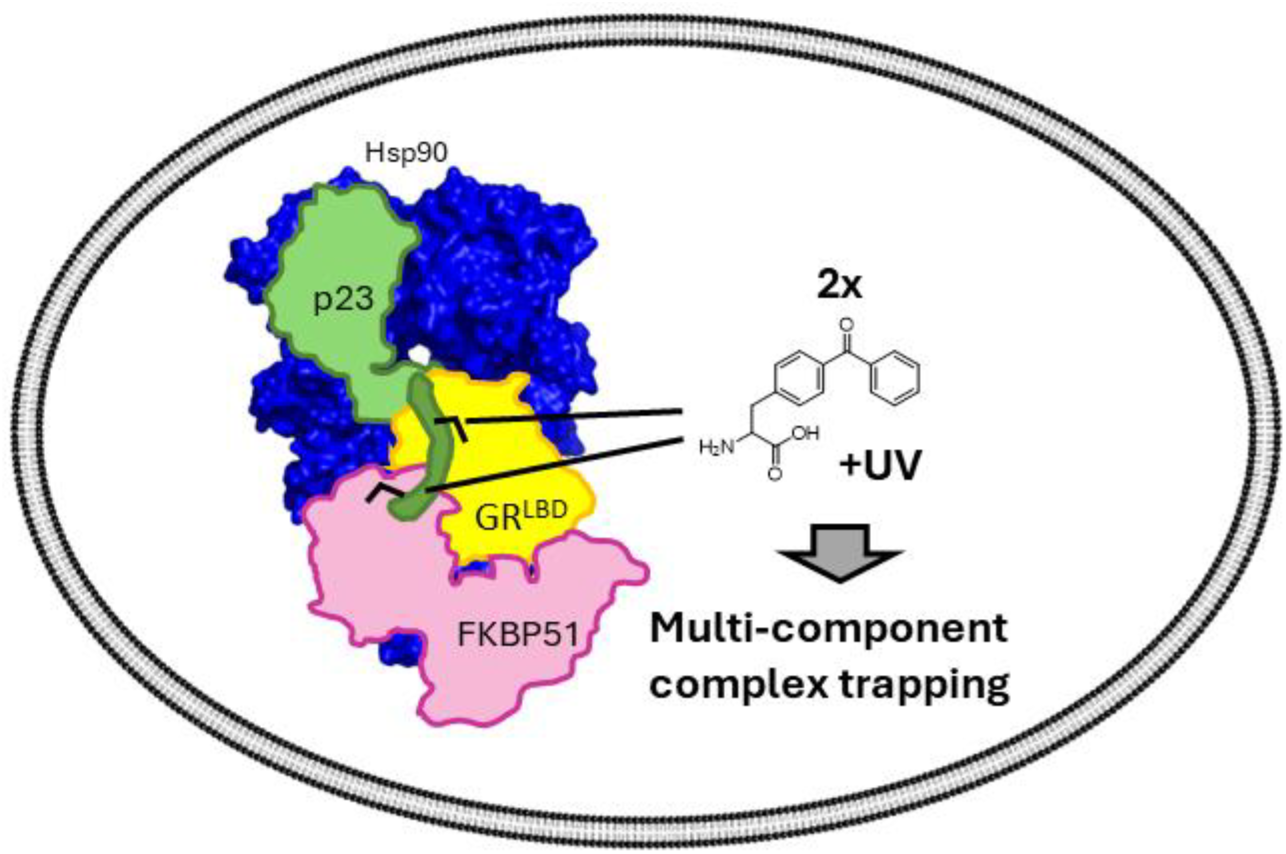

## Introduction

The glucocorticoid receptor (GR) is a member of the steroid hormone receptor (SHR) family and is crucial for key endocrine processes including stress hormone signaling and regulation of inflammation [1]. Prior to activation, the GR undergoes a complex maturation cycle that critically depends on the heat shock protein 90 (Hsp90) machinery and is aided in its final steps by co-chaperones such as FK506 binding proteins 51 and 52 (FKBP51 and FKBP52, respectively) and p23 [1], [2].

p23 was found to bind to the Hsp90 dimer during client maturation [3] and is essential for GR activity [4], [5]. Inhibition or delaying of ATPase function of Hsp90 has been suggested as one way in which p23 promotes GR signaling. This would keep the GR tethered to Hsp90, thereby prolonging the time for correct folding of the GR [6]. However, overexpression of p23 was also found to reduce GR signaling, possibly by attenuating the release of the activated GR from the Hsp90 complex [7]. Recently, p23 was shown to directly interact with GR via its C-terminal tail helix (amino acids 117-131) at the beginning of the intrinsically disordered C-terminal domain [4], [5], [8].

The co-chaperone FKBP51 is known for its inhibitory activity on the GR whereas FKBP52 promotes GR signaling [9], [10], [11], [12]. FKBP51 plays a key role in the control of mammalian stress response and has emerged as a promising target for stress-related disorders [12], [13], [14].

The GR is the most studied Hsp90 substrate and serves as a model client for the Hsp90 machinery [4], [12], [15], [16]. However, the interplay between the different co-chaperones is poorly understood. In particular, the mechanism through which FKBP51 inhibits the GR and the molecular basis of the opposing effects of FKBP51 vs FKBP52 have largely remained elusive. In this paper, we provide evidence for the existence of an FKBP51-GR-p23-Hsp90_2_ multi-cochaperone complex which is exclusive to FKBP51 vs FKBP52, representing an important state of apo-GR in living human cells and providing a refined mechanism for regulating GR signaling in human cells.

## Results and Discussion

### The GR has a well-defined p23 interaction site *in cellulo*

We previously investigated the interaction of GR with its co-chaperones FKBP51 and FKBP52 via photoinducible crosslinking [12]. To this end, we utilized amber suppression to incorporate the unnatural amino acid para-benzoyl phenylalanine (pBpa) in a series of experiments into defined positions on the surface of GR. Upon UV irradiation, pBpa can form covalent photocrosslinks with other amino acids in the vicinity. After cell lysis, proteins that were directly contacting the probed position can be detected by a gel shift in Western Blots.

To analyze the p23-contacting surface of GR in human cells, we probed a collection of GR pBpa mutants covering most surface amino acids of the ligand binding domain of GR (GR^LBD^) and investigated the interaction with p23 via Western Blotting (Supplementary Figures 1 and 2) [12]. This revealed a well-defined interaction site of GR with p23 on one side of the GR^LBD^, mainly around helix 9 and 10 of GR. These residues fit remarkably well with the cryo-EM structure of GR-Hsp90_2_-p23, which showed an interaction of p23^117-131^, a helical subdomain structure, with the hydrophobic grove of helix 9 and 10 (Figure 1 B and Supplementary Figure 2) [15]. However, unlike the cryo-EM structure which was obtained with the ligand-bound, activated GR, our findings reflect the unliganded apoGR since the crosslinks were disrupted by pre-treatment with the GR agonist dexamethasone [12].

**Figure 1:**
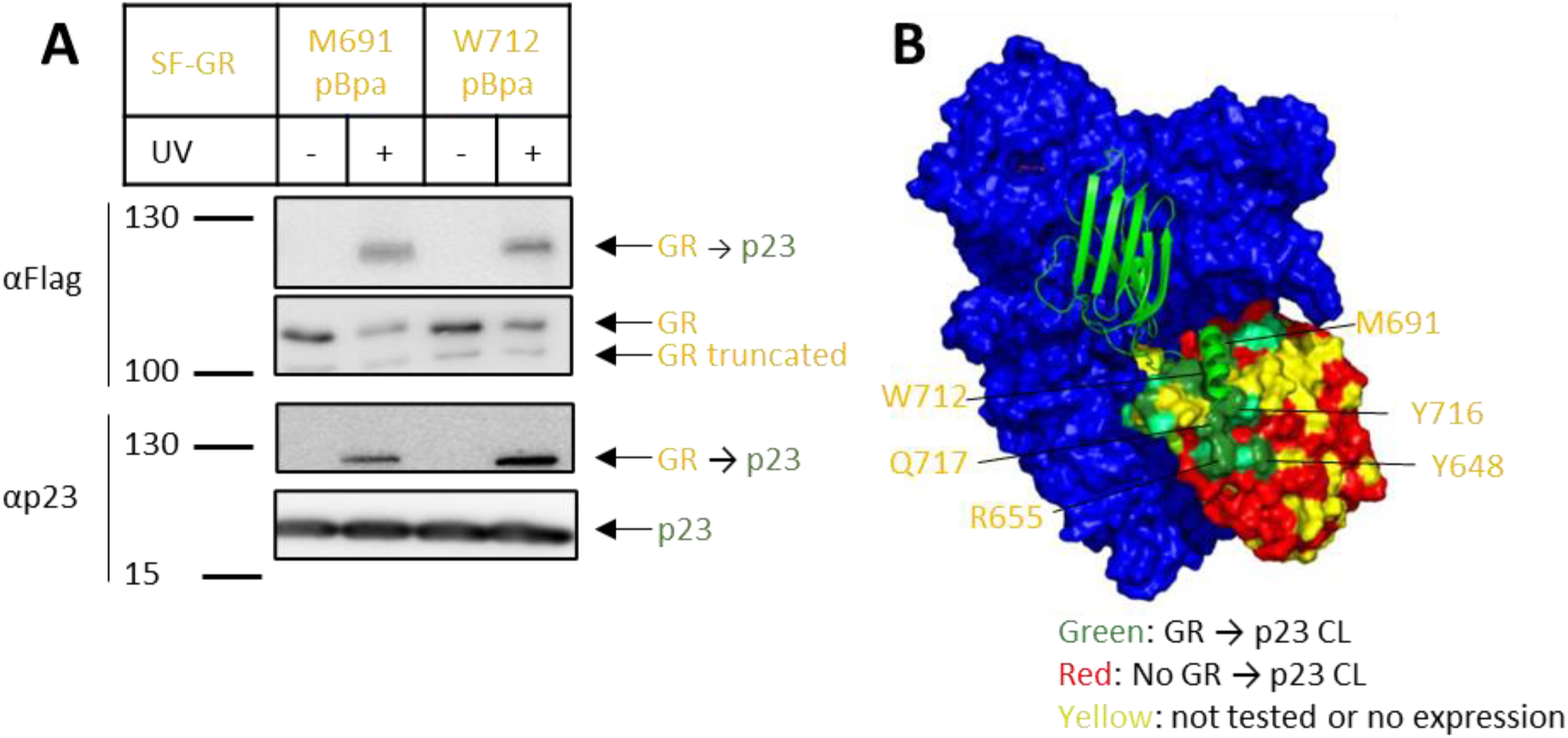
In-cell site-specific photocrosslinking reveals a well-defined GR-p23 interaction site: **A:** HEK293 cells overexpressing representative Flag-tagged GR pBpa mutants were UV irradiated for 30 or 0 minutes and analyzed for GR→p23 crosslinks by Western Blot. Apparent molecular weight (in kDa) and primary antibodies are indicated on the left and protein identities are annotated on the right of the blot. Bands at approximately 120 kDa are indicative of a GR→p23 crosslink. **B**: Positions tested by in-cell photocrosslinking were mapped on the structure of the GR-Hsp90_2_-p23 complex (PDB: 7KRJ, GR with green/yellow/red surface, p23 in green cartoon, Hsp90_2_ surface in blue). Positions tested positive for GR→p23 crosslinks are shown in green (respective amino acids are annotated on human GR), positions negative for GR→p23 crosslinks are shown in red. Positions which did not express as pBpa mutants or which were not tested are shown in yellow. Primary data is shown Supplementary Figures 1 and 2.

### FKBP51 stabilizes the GR-p23 interaction *in cellulo*

We next explored the interplay between the co-chaperones p23, FKBP51 and FKBP52 within the Hsp90-GR heterocomplex. In recently solved cryo-EM structures of the FKBP51-Hsp90_2_-GR and FKBP52-Hsp90_2_-GR complexes [15], the GR adopts a strikingly different binding mode compared to the p23-Hsp90_2_-GR complex [4], with the p23 binding site on the GR around M691, W712 and Q717 being masked by Hsp90. We thus expected FKBP51 and FKBP52 to compete with the GR-p23 interaction.

To our great surprise, we observed that FKBP51 significantly increases the intensity of the GR-p23 crosslink at three investigated positions while FKBP52 does not (Figure 2 A - F). Similar results were observed for all other GR^LBD^ positions known to crosslink to p23 with enhanced GR→p23 crosslink intensity after FKBP51 co-overexpression and unchanged or decreased GR→p23 crosslink intensity after co-overexpressing FKBP52 (Supplementary Figure 3). This revealed a strengthening of the GR-p23 interaction in the presence of FKBP51, while FKBP52 weakens the p23-GR interaction.

**Figure 2:**
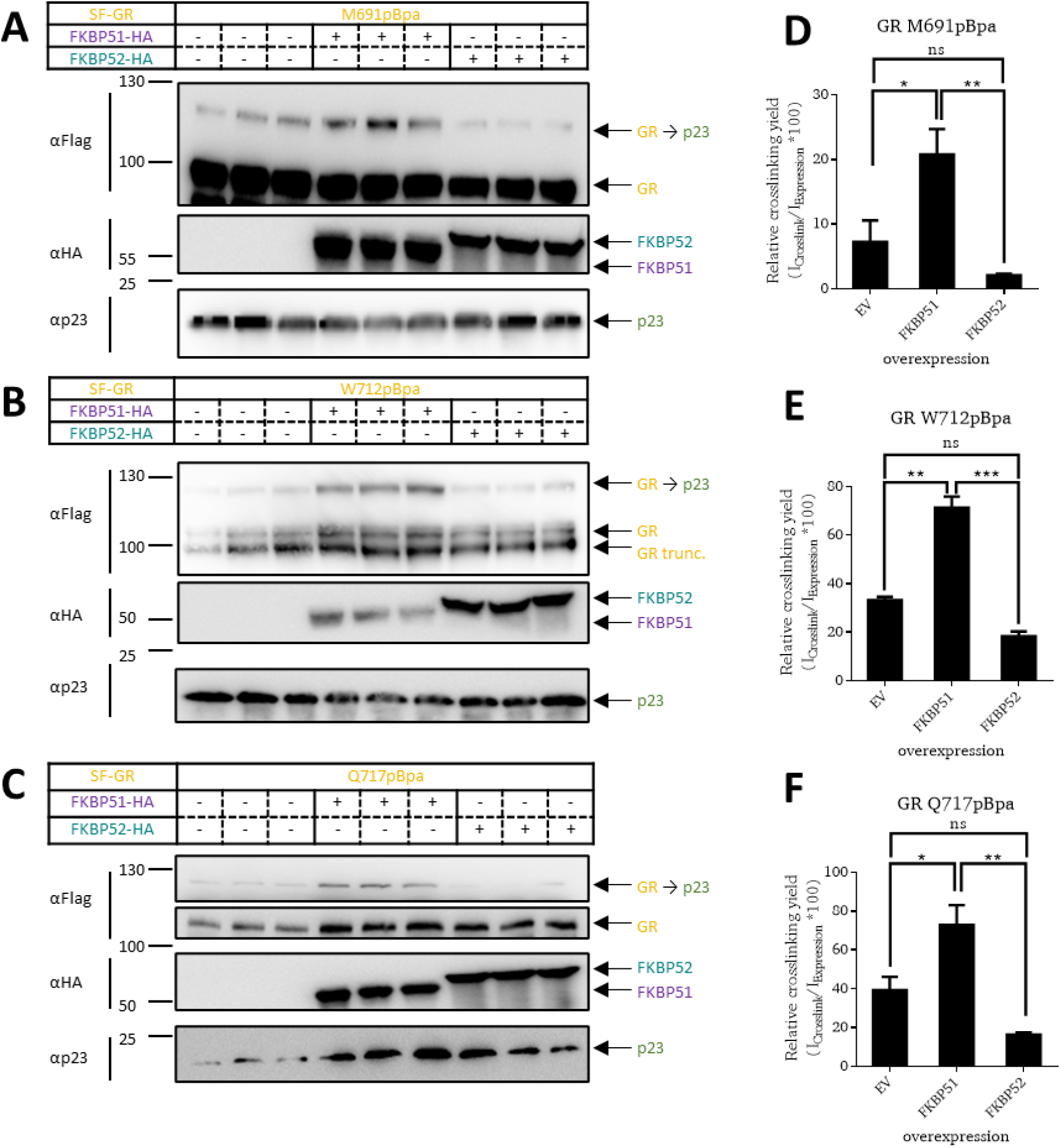
FKBP51 stabilizes the GR-p23 interaction in cellulo: **A – C**: HEK293 cells co-overexpressing Flag-tagged GR pBpa mutants and HA-tagged FKBP51 or FKBP52 were UV irradiated and GR→p23 crosslinks were analyzed by Western Blot in biological triplicates. Apparent molecular weight (in kDa) and primary antibodies are indicated on the left and protein identities are annotated on the right of the blot. **D – F:** Quantification of the biological triplicates of GR→p23 crosslinking bands, normalized to the GR expression bands, are shown to the right. Error in standard deviation.

Position 119 in the proline-rich loop of FKBP51 and FKBP52 has been suggested to be important for GR regulation and a determinant of the opposing effects of FKBP51 and FKBP52. Replacing Leu119 in FKBP51 with proline diminished the inhibiting effect of FKBP51 on the GR, while the reverse swap in FKBP52 (Pro119 to Leu119) decreased its activating effect on GR [17]. To refine the molecular basis for the GR-p23 stabilization by FKBP51 but not FKBP52, we tested the FKBP51^L119P^ and FKBP52^P119L^ mutants for in the GR->p23 in-cell photocrosslinking model (Figure 3 and Supplementary Figure 4). FKBP51^L119P^ was substantially less capable of GR->p23 crosslinking compared to wildtype FKBP51. Conversely, FKBP52^P119L^ overexpression, the GR->p23 crosslink intensity increased slightly compared to wildtype FKBP52.

**Figure 3:**
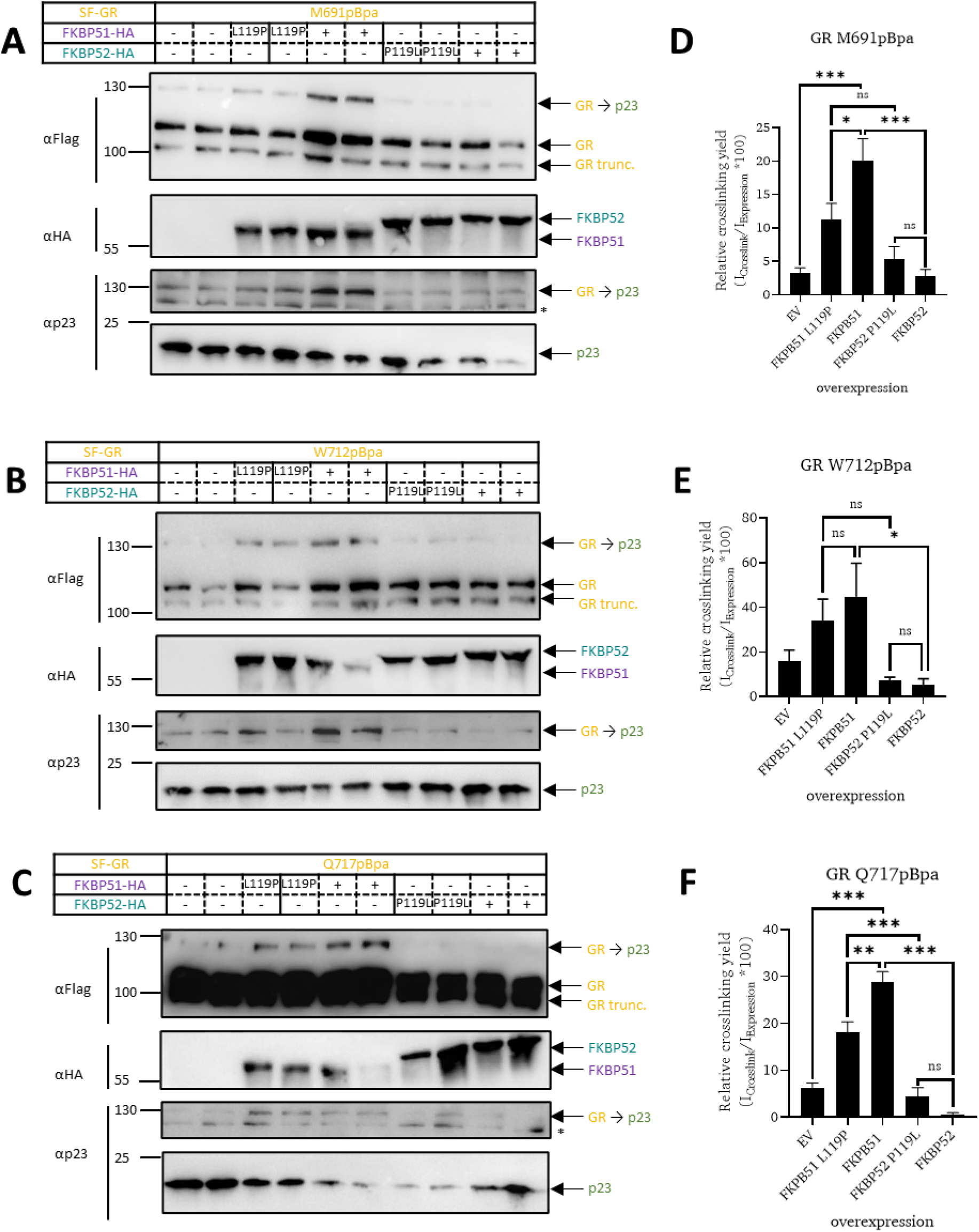
Amino acid swap at position 119 reduces the stabilizing effect of FKBP51 on the GR-p23 interaction in cellulo: **A:** HEK293 cells co-overexpressing Flag-tagged GR pBpa mutants and HA-tagged FKBP51, FKBP51^L119P^, FKBP52^P119L^ or FKBP52 were UV irradiated and GR→p23 crosslinks were analyzed by Western Blot (representative biological duplicates shown). Protein sizes and primary antibodies are indicated on the left and protein identities are annotated on the right of the blot. **B:** Quantification of the six biological replicates (visualized in 3 different blots, see Supplementary Figure 4 of GR→p23 crosslinking bands, normalized to the StrepFlag-GR expression bands, are shown to the right. EV=empty vector control, error bars represent standard error, star marks unspecific bands.

### FKBP51 and p23 can bind to the GR simultaneously

We suspected that FKBP51 stabilizes the GR-p23 interaction by binding them in a higher-order complex. To further investigate how FKBP51 influences the GR-p23 interaction, we performed crosslinking of all three proteins simultaneously. By expressing multiple proteins with orthogonal tags and a pBpa incorporation site in two proteins, the architecture of multi-component complexes can be analyzed (Figure 4).

**Figure 4:**
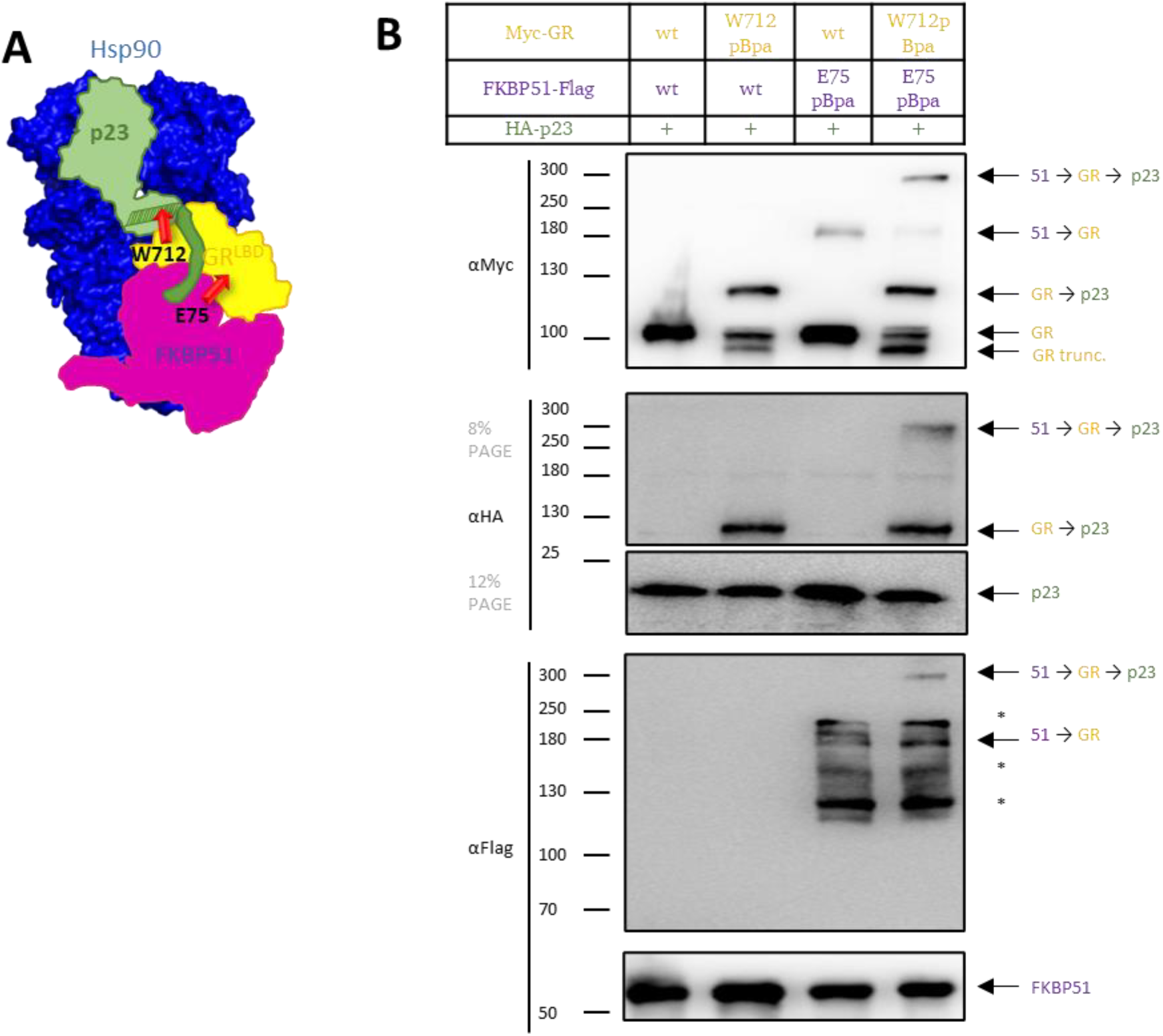
FKBP51 and p23 can bind to the GR simultaneously, analyzed by double crosslinks: **A**: Scheme of the chain-crosslinked complex. FKBP51^E75pBpa^ crosslinks to GR, GR^W712pBpa^ crosslinks to p23. **B:** HEK293 cells co-overexpressing Myc-tagged wildtype GR or GR^W712pBpa^, Flag-tagged wildtype FKBP51 or FKBP51^E75pBpa^ and HA-tagged p23 were UV irradiated and the photocrosslinks were analyzed by Western Blot. Apparent molecular weight (in kDa) and protein identities are indicated left and right of the blot respectively; stars mark unspecific bands. Photocrosslinked bands at approximately 300 kDa correspond to the postulated FKBP51→GR→p23 multi-chaperone complex.

Specifically, we used FKBP51^E75pBpa^ known to crosslink to the GR and GR^W712pBpa^ known to crosslink to p23 (Figure 4 A). After co-overexpressing these mutants together with HA-tagged p23, we observed the existence of a larger complex which contains FKBP51, GR and p23 (Figure 4 B). The formation of this complex strictly depended on the presence of both pBpa mutants. Notably, the complex consisting of FKBP51-GR-p23 seems to be favored over a complex only consisting of FKBP51 and the GR, reflected by a stronger band for the FKBP51^E75pBpa^→GR^W712pBpa^→p23 complex compared to the FKBP51^E75pBpa^→GR^W712pBpa^ band. Repeating the same experiment with FKBP52^D75pBpa^ revealed no evidence for a FKBP52-GR-p23 complex (Supplementary Figure 5). FKBP51 might enhance the GR-p23 interaction by stabilizing the GR in a p23 binding-competent conformation or by direct binding to p23.

### The C-terminal tail of p23 contacts FKBP51 in the presence of the GR

To investigate whether FKBP51 stabilizes the GR in a p23 binding competent state or directly binds to p23 in the complex, we screened various FKBP51 pBpa mutants for photoinducible crosslinking to p23 (Supplementary Figure 6). We found several positions in the FK1 domain of FKBP51 that directly crosslink to p23 in dependence of Hsp90 (Supplementary Figure 6). Intriguingly, some of these positions were close or even overlapped with the GR binding surface of FKBP51 (Figure 5 A and B). To explore this further, we repeated the analysis of FKBP51→p23 crosslinks with and without GR co-overexpression. Several positions such as FKBP51^N63pBpa^ were unaffected by GR overexpression (Figure 5 D). However, other p23 contacts (especially in the proline-rich loop overhanging the FK506-binding site) were clearly abolished in the presence of GR (Figure 5 F). These latter positions such as FKBP51^P123pBpa^ switch from contacting p23 to contacting GR if the latter is available. This is reversible as the GR-sensitive FKBP51→p23 crosslinks reappeared after dexamethasone treatment, which leads to dissociation of GR from the Hsp90 complex.

**Figure 5:**
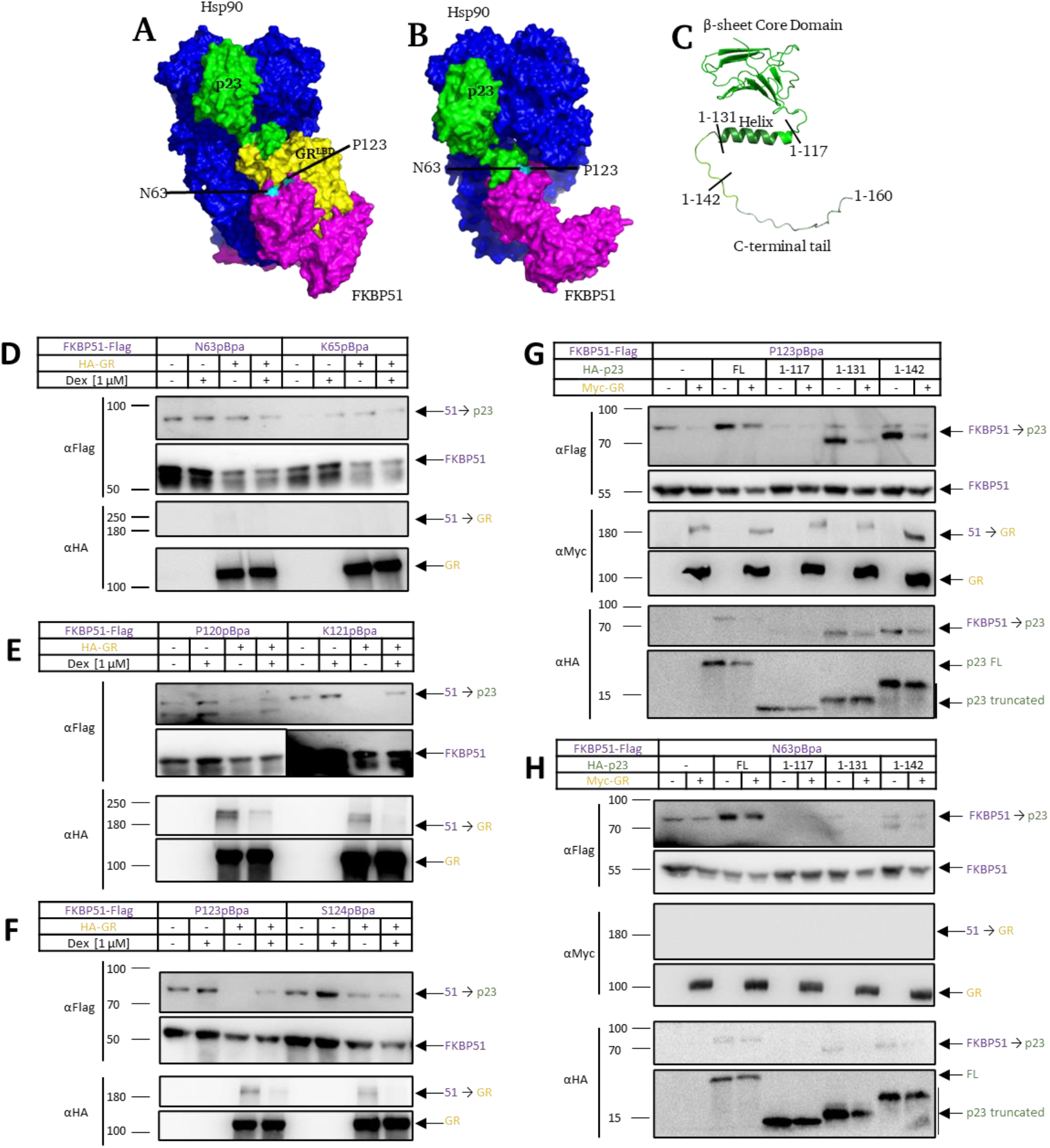
FKBP51 interacts with the C-terminal tail of p23 in presence of the GR: A and B: Schematic representation of possible FKBP51-p23 complexes with (**A**, FKBP51 from 8FFW merged with GR-p23 from 7KRJ) and without GR (**B**, FKBP51 from 7L7I merged with p23 from 7KRJ, both shown on the same side of Hsp90_2_). p23 is resolved up to amino acid D133. FKBP51^N63^ and ^P123^ are marked in cyan and teal, respectively. **C**: Scheme of p23 truncation constructs shown for the AlphaFold2 model of p23 (AF-Q15185-F1). **<u>D – F:</u>** HEK293 cells overexpressing Flag-tagged FKBP51 pBpa mutants with and without co-overexpression of HA-tagged GR were optionally pre-treated with Dexamethasone, UV irradiated, and FKBP51→p23 crosslinks were analyzed by Western Blot. **G and H:** HEK293 cells co-overexpressing Flag-tagged FKBP5^P123pBpa^ or FKBP51^N63pBpa^, full length or truncated HA-tagged p23 variants and optionally Myc-tagged GR were UV irradiated and the crosslinks of FKBP51→p23 were analyzed by Western Blot. Protein sizes and protein identities are indicated left and right of the blot, respectively.

The persistence of certain FKBP51→p23 crosslinks in the presence of GR indicates that p23 can still directly contact FKBP51 in the FKBP51-Hsp90_2_-GR-p23 multicomponent complex. To refine these FKBP51-p23 interactions, we mapped the role of the C-terminal tail of p23, which is unresolved in the cryo-EM structures but could reach out to the putative position on the FKBP51 FK1 domain such as N63 (Figure 5 B). We therefore tested p23 truncation variants missing parts of the C-terminal tail (Figure 5 C, [18]) for their crosslinking behavior to FKBP51. The crosslinks of FKBP51^P123pBpa^ to endogenous p23 are clearly suppressed by co-overexpressing truncated variants of p23, indicating an efficient intracellular competition of all overexpressed p23 variants with endogenous p23 in the Hsp90 complex (Figure 5 G). In the absence of GR, FKBP51^P123pBpa^ efficiently crosslinked to p23^1-142^ and p23^1-131^ but not p23^1-117^, indicating binding of the proline-rich loop of FKBP51 to amino acids 118 to 131 of p23. Conversely, FKBP51^N63pBpa^ only crosslinked to the full length p23, but not p23^1-142^ or shorter variants, suggesting an interaction with the C-terminal tail of p23 (amino acids 142 to 160). The FKBP51^P123Bpa^ crosslinks to the truncated p23^1-131^ and p23^1-142^ constructs were both disrupted by GR co-overexpression, as observed before for full-length p23. In contrast to FKBP51^P123Bpa^, FKBP51^N63Bpa^ very poorly crosslinked to p23^1-131^ and p23^1-142^, especially in the presence of GR. The C-terminus (142-160) thus seems to be a major interaction site with FKBP51 in the FKBP51-Hsp90_2_-GR-p23 multicomponent complex (Figure 5 H).

## Discussion

Efficient GR hormone binding relies on a precisely orchestrated interplay with the Hsp90- (co)chaperone machinery. Co-chaperones are crucial to fine-tune physiologically adequate GR activation. This is most apparent for the antagonistic effects of FKBP51 and FKBP52 and the pathological consequences of excessive FKBP51 overexpression. Recent advancements in cryo-EM structures and structural elucidation by photoinducible in-cell crosslinking shed light on the interaction of the GR with its co-chaperones FKBP51, FKBP52 and p23 [4], [12], [15], but the functional implication for the GR regulation have been less clear.

Utilizing photoinducible crosslinking, we confirmed the major GR^LBD^-p23 interaction site *in cellulo*. In accordance with the cryo-EM structure, we observed a well-defined interaction site located at helices 9 and 10 of GR. The cryo-EM studies observed a dexamethasone-bound state trapped in a state before dissociation while our studies represent the apoGR. This indicates a strong interaction between GR and the C-terminal helix of p23 that is present before and after GR activation and might be only lost after the dissociation of p23 from the GR-Hsp90_2_ complex.

FKBP51 and FKBP52 are key players in GR regulation with opposing roles despite their high structural similarity and a surprisingly similar overall interaction pattern with GR [12]. Analyzing the interplay between these co-chaperones, we found that FKBP51 stabilizes the GR-p23 interaction while FKBP52 does not. We also show that FKBP51 and p23 are part of the same GR-Hsp90_2_ complex. Steroid hormone-Hsp90_2_ complexes that contain both p23 and FKBP51 are consistent with previous observations from co-immunoprecipitations and native MS studies [9], [16], [19], [20] but are incompatible with the cryo-EM structures of FKBP51-Hsp90_2_-GR [15].

The closed form of the Hsp90 dimer is the preferred state for binding of p23 and FKBP51. Previous studies suggested that FKBP51 and p23 reside on opposite sides of the Hsp90 dimer [21], while GR^LBD^ and p23 as well as GR^LBD^ and FKBP51 are positioned on the same side of the Hsp90 dimer. Utilizing in-cell photocrosslinking, we found evidence for an FKBP51-Hsp90_2_-GR-p23 complex where FKBP51 and p23 not only are located on the same side of the Hsp90 dimer but are in direct contact with each other in the presence of GR (Figure 4 and Figure 5). More specifically, p23 contacts the FK1 domain of FKBP51 with its C-terminal tail. At this interaction interface, FKBP51 but not FKBP52 seems to provide an additional binding surface for p23 and thereby stabilizing p23 in the complex.

Recent cryo-EM data showed two dramatically different configurations of the GR^LBD^ in the Hsp90 complex when bound to p23 [4] or FKBPs [15]. One key feature in the former structure is the GR-p23 interaction site provided by the hydrophobic patch at helix 9 and 10 of GR. This interaction site is not available for p23 binding in the structures bound with either FKBP51 or FKBP52 where GR is rotated and this patch is masked by Hsp90, implying exclusive binding of either FKBPs or p23 (Figure 6 A). However, on the basis of our data, generated by *in cellulo* crosslinking, we propose that FKBP51 preferentially interacts with the GR-p23 complex while FKBP52 does not (Figure 6 B). This suggests that the configuration of the GR-p23 complex is more compatible with FKBP51 binding while FKBP52 prefers binding to the rotated GR^LBD^.

**Figure 6:**
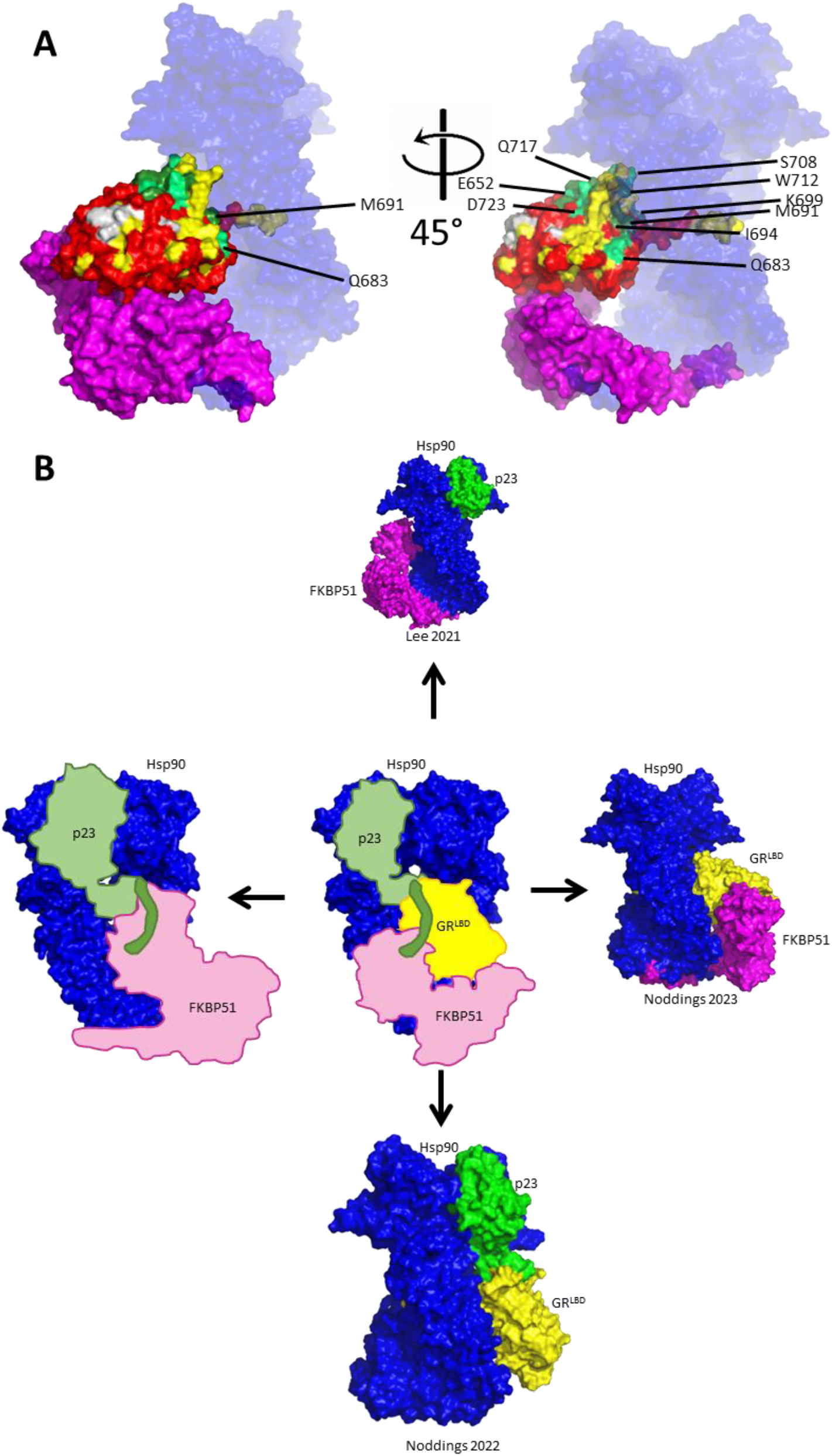
**A: FKBP51→GR→p23 crosslinks are not compatible with a cryo-EM structure of the FKBP51-GR-Hsp90_2_ complex:** GR→p23 crosslinks superimposed on GR in complex with FKBP51 (PDB: 8FFW, FKBP51 shown in magenta, Hsp90_2_ shown in transparent blue for better visibility). Positions positive for GR→p23 crosslinks are shown in green (light green: weak crosslinks, dark green: strong crosslink), positions negative for GR→p23 crosslinks are shown in red, GR positions not assessed are shown in yellow. **B: Architecture of a novel FKBP51-Hsp90**_**2**_**-GR-p23 complex**. Comparison of proposed multi-co-chaperone complex with experimentally observed GR-p23-Hsp90_2_, FKBP51-GR -Hsp90_2_ or FKBP51-p23-Hsp90_2_ complexes. The original publications are indicated below each complex.

In the FKBP51-Hsp90_2_-GR-p23 complex, we suggest that FKBP51 binds to GR with its proline-rich loop (that includes L119) and to the C-terminal loop of p23. However, we cannot exclude the possibility that FKBP51 binds to other parts of p23 by interactions that are indirectly stabilized by the C-terminal tail of p23. Likewise, the amino acid in position 119 might affect the interaction with GR or p23 allosterically. In contrast to recently published *in vitro* data, we observe a strong interaction of the FK1 domain of FKBP51 with p23 which might reflect differences between in-cell conditions and isolated purified proteins [22].

Our data suggests a refined model for the inhibitory effect of FKBP51 on GR activity. FKBP51 stabilizes p23 in complex with Hsp90_2_-GR. By slowing p23 dissociation, FKBP51 disfavors disassembly of the Hsp90_2_-GR complex, which is thought to be necessary for GR to act as a transcription factor, thereby attenuating GR signaling.

On the other hand, FKBP52 does not stabilize p23 or even disfavors p23 in the complex and thus promotes Hsp90_2_-GR disassembly.

One reason for the opposing effects of FKBP51 and FKBP52 on the GR-p23 interaction might be the difference in surface amino acids of FKBP51 and FKBP52. Notably, the FK1 domain of FKBP51 is more positively charged compared to FKBP52. The former might therefore interact more efficiently with the C-terminus of p23, which is highly negatively charged. The proline rich loop of FKBP51 has been discussed as one of the key elements of FKBP51 necessary for its co-chaperone activity [17]. By swapping the key amino acid at position 119 between FKBP51 and FKBP52, we could demonstrate an impact on the stabilization of p23 in the GR-Hsp90 complex.

Taken together, we provide evidence for a FKBP51-Hsp90_2_-GR-p23 complex and a mechanistic explanation for the functional differences between FKBP51 and FKBP52 with regards to their impact on GR activity.

## Material and Methods

### Cell culture

HEK293 cells were cultivated at 37 °C and 5% CO_2_ in DMEM supplemented with 10% FBS and 1% penicillin-streptomycin solution.

### Amber Suppression

As described previously by Baischew et al. 2023. In brief: 200 000 HEK293 cells were seeded in a Poly-L-Lysine coated 12 well plate and were grown over night to approximately 60% confluency. Cells were transfected using PEI Prime transfection reagent. For each well of a 12 well plate, 1.5 μg PEI, 300 ng Synthetase (pBpa tRNA Synthetase p_NEU_EBpaRS_4xBstYam with four copies of U6-BstYam expression cassettes[23], gift from I. Coin), 300 ng GR TAG mutant, 200 ng FKBP51 TAG mutant and 100 ng HA-p23 were added to 100 μl OptiMEM and incubated for 30 minutes. The medium was replaced with fresh medium containing 500 μM pBpa (50 mM in 100 mM NaOH, sterile filtered) and the transfection mix was added. The cells were incubated for approximately 40 hours to allow for protein expression and UV irradiated (2 × 15 W, Conso VL-215.L) for 30 minutes on ice.

For FKBP51/FKBP52 switch mutants, the same protocol was used but with 150 000 HEK293 cells, 400 ng pBpa tRNA Synthetase, 400 ng GR TAG mutant, 300 ng FKBP51^L119p^ or 400 ng FKBP52^P119L^ per well.

### Immunoblotting

Cells were lysed in 100 μl NETN lysis buffer (100 mM NaCl, 20 mM Tris pH 8, 0,5 mM EDTA, 0.5% Nonidet P-40, protease inhibitor cocktail (Roche)) for 30 minutes on ice while shaking. The lysate was collected in a reaction tube and centrifuged for 15 minutes at 15 000 g, 4 °C. The supernatant was transferred to a new reaction tube and supplemented with 4x Laemmli buffer and boiled for 10 minutes. The proteins were separated by SDS-PAGE and transferred to a nitrocellulose membrane (Ambersham). The membrane was blocked for at least 30 minutes in 5% milk in TBS and incubated overnight at 4 °C under movement with the primary antibody. After incubation, the membranes were washed 3×5 minutes with TBS and, if necessary, incubate for at least one hour with an appropriate secondary, HRP-coupled antibody diluted in 5% milk in TBS. Excess secondary antibody was washed off 3×5 minutes with TBS and the band intensity was measured with an image analyzer (Fuji Photo Film).

The following antibodies were used: Anti-Flag (mouse monoclonal clone M2, Sigma-Aldrich) 1:5,000; anti-HA (rat monoclonal, clone 3F10, Roche) 1:1,000; anti-p23 (mouse monoclonal, clone JJ6, Santa Cruz Biotechnology (SCBT), anti-Myc (2276S, CST) 1:5000 in 5% BSA in TBS.

### Statistical Analysis

Statistical analysis of Western blots were performed by one-way ANOWA. Significances are donated as follows: n.s. P > 0.05; * P ≤ 0.05; **P ≤ 0.01; ***P ≤ 0.001

### Cloning

Golden Gate cloning of TAG mutants was performed as described previously by Baischew et al. 2023.

