## Supplementary Figures for "A new FKBP51-GR-p23-Hsp90_2_ multi-cochaperone complex identified and characterized by site-specific in-cell photocrosslinking"

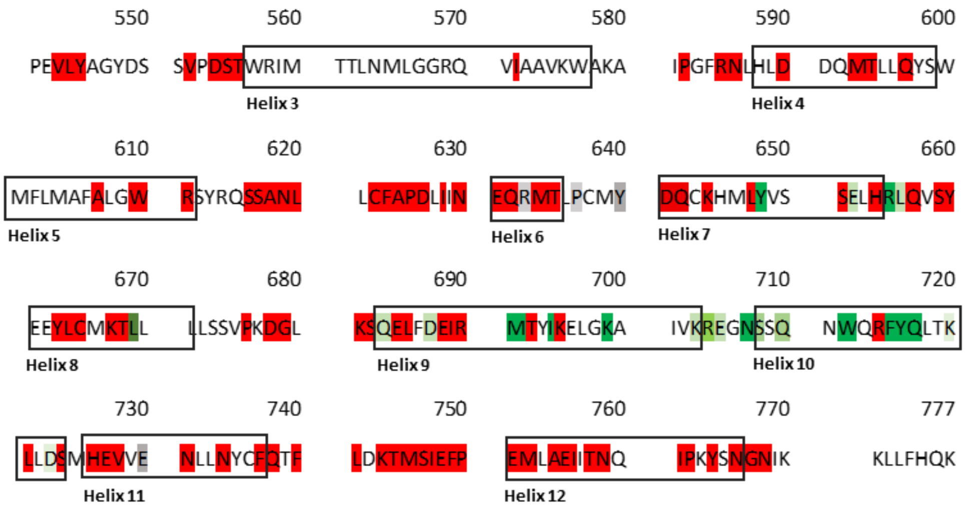


*Supplemental Figure 1: **Result of GR→p23 screening**: Positions which were tested positive for GR→p23 crosslink are marked in green (once: lime green, twice: chartreuse, three times: dark green), positions negative for GR→p23 crosslink are marked in red. Positions which did not show an expression band are marked in grey. Helixes in LBD are labeled.*

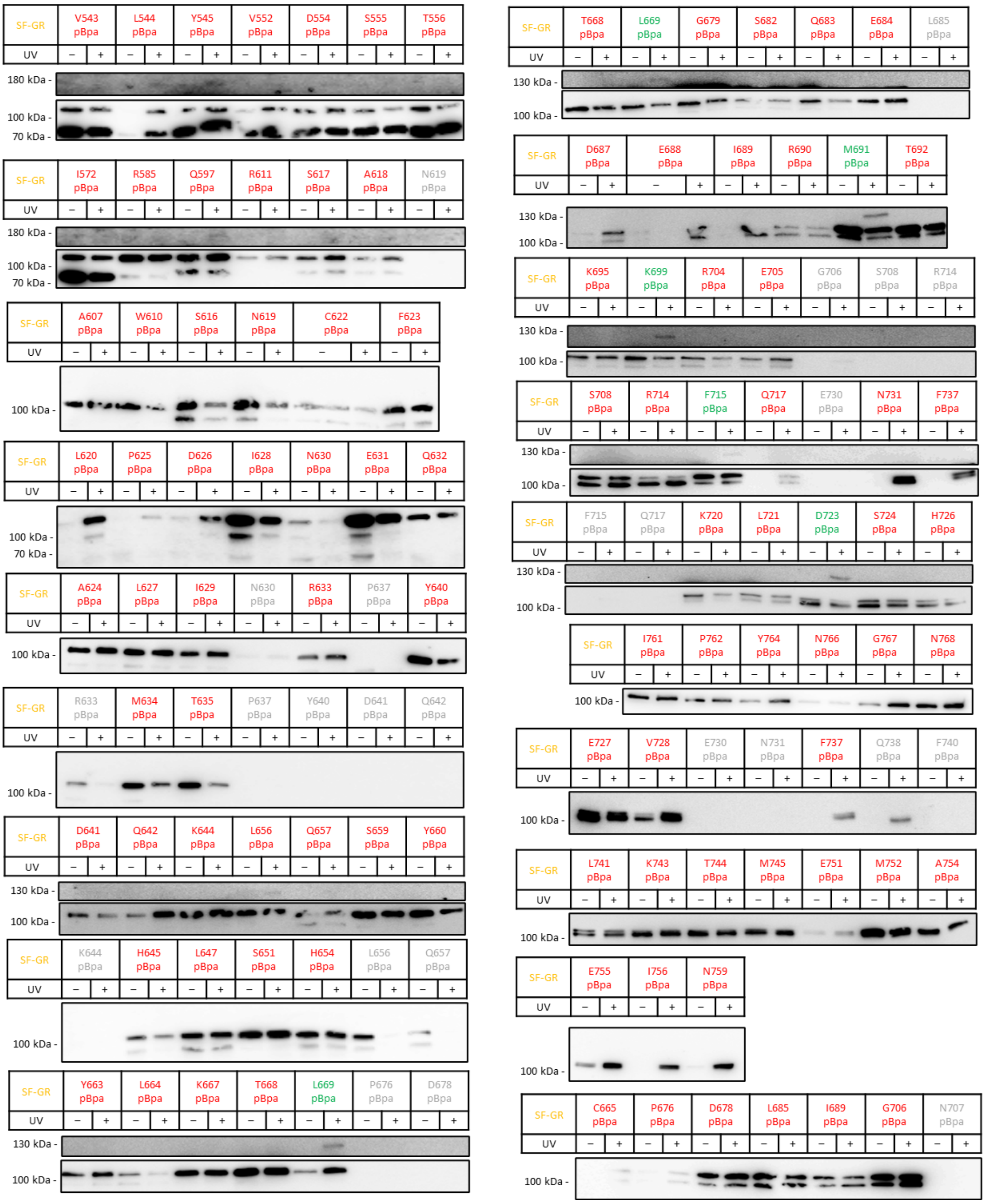

SF-GR V543 pBpa L544 pBpa Y545 pBpa V552 pBpa D554 pBpa S555 pBpa T556 pBpa
UV – + – + – + – + – + – + – +
180 kDa -
100 kDa -
70 kDa -
SF-GR I572 pBpa R585 pBpa Q597 pBpa R611 pBpa S617 pBpa A618 pBpa N619 pBpa
UV – + – + – + – + – + – + – +
180 kDa -
100 kDa -
70 kDa -
SF-GR A607 pBpa W610 pBpa S616 pBpa N619 pBpa C622 pBpa F623 pBpa
UV – + – + – + – + – + – +
100 kDa -
SF-GR L620 pBpa P625 pBpa D626 pBpa I628 pBpa N630 pBpa E631 pBpa Q632 pBpa
UV – + – + – + – + – + – + – +
100 kDa -
70 kDa -
SF-GR A624 pBpa L627 pBpa I629 pBpa N630 pBpa R633 pBpa P637 pBpa Y640 pBpa
UV – + – + – + – + – + – + – +
100 kDa -
SF-GR R633 pBpa M634 pBpa T635 pBpa P637 pBpa Y640 pBpa D641 pBpa Q642 pBpa
UV – + – + – + – + – + – + – +
100 kDa -
SF-GR D641 pBpa Q642 pBpa K644 pBpa L656 pBpa Q657 pBpa S659 pBpa Y660 pBpa
UV – + – + – + – + – + – + – +
130 kDa -
100 kDa -
SF-GR K644 pBpa H645 pBpa L647 pBpa S651 pBpa H654 pBpa L656 pBpa Q657 pBpa
UV – + – + – + – + – + – + – +
100 kDa -
SF-GR Y663 pBpa L664 pBpa K667 pBpa T668 pBpa L669 pBpa P676 pBpa D678 pBpa
UV – + – + – + – + – + – + – +
130 kDa -
100 kDa -
SF-GR T668 pBpa L669 pBpa G679 pBpa S682 pBpa Q683 pBpa E684 pBpa L685 pBpa
UV – + – + – + – + – + – + – +
130 kDa -
100 kDa -
SF-GR D687 pBpa E688 pBpa I689 pBpa R690 pBpa M691 pBpa T692 pBpa
UV – + – + – + – + – + – +
130 kDa -
100 kDa -
SF-GR K695 pBpa K699 pBpa R704 pBpa E705 pBpa G706 pBpa S708 pBpa R714 pBpa
UV – + – + – + – + – + – + – +
130 kDa -
100 kDa -
SF-GR S708 pBpa R714 pBpa F715 pBpa Q717 pBpa E730 pBpa N731 pBpa F737 pBpa
UV – + – + – + – + – + – + – +
130 kDa -
100 kDa -
SF-GR F715 pBpa Q717 pBpa K720 pBpa L721 pBpa D723 pBpa S724 pBpa H726 pBpa
UV – + – + – + – + – + – + – +
130 kDa -
100 kDa -
SF-GR I761 pBpa P762 pBpa Y764 pBpa N766 pBpa G767 pBpa N768 pBpa
UV – + – + – + – + – + – +
100 kDa -
SF-GR E727 pBpa V728 pBpa E730 pBpa N731 pBpa F737 pBpa Q738 pBpa F740 pBpa
UV – + – + – + – + – + – + – +
100 kDa -
SF-GR L741 pBpa K743 pBpa T744 pBpa M745 pBpa E751 pBpa M752 pBpa A754 pBpa
UV – + – + – + – + – + – + – +
100 kDa -
SF-GR E755 pBpa I756 pBpa N759 pBpa
UV – + – + – +
100 kDa -
SF-GR C665 pBpa P676 pBpa D678 pBpa L685 pBpa I689 pBpa G706 pBpa N707 pBpa
UV – + – + – + – + – + – + – +
100 kDa -


21

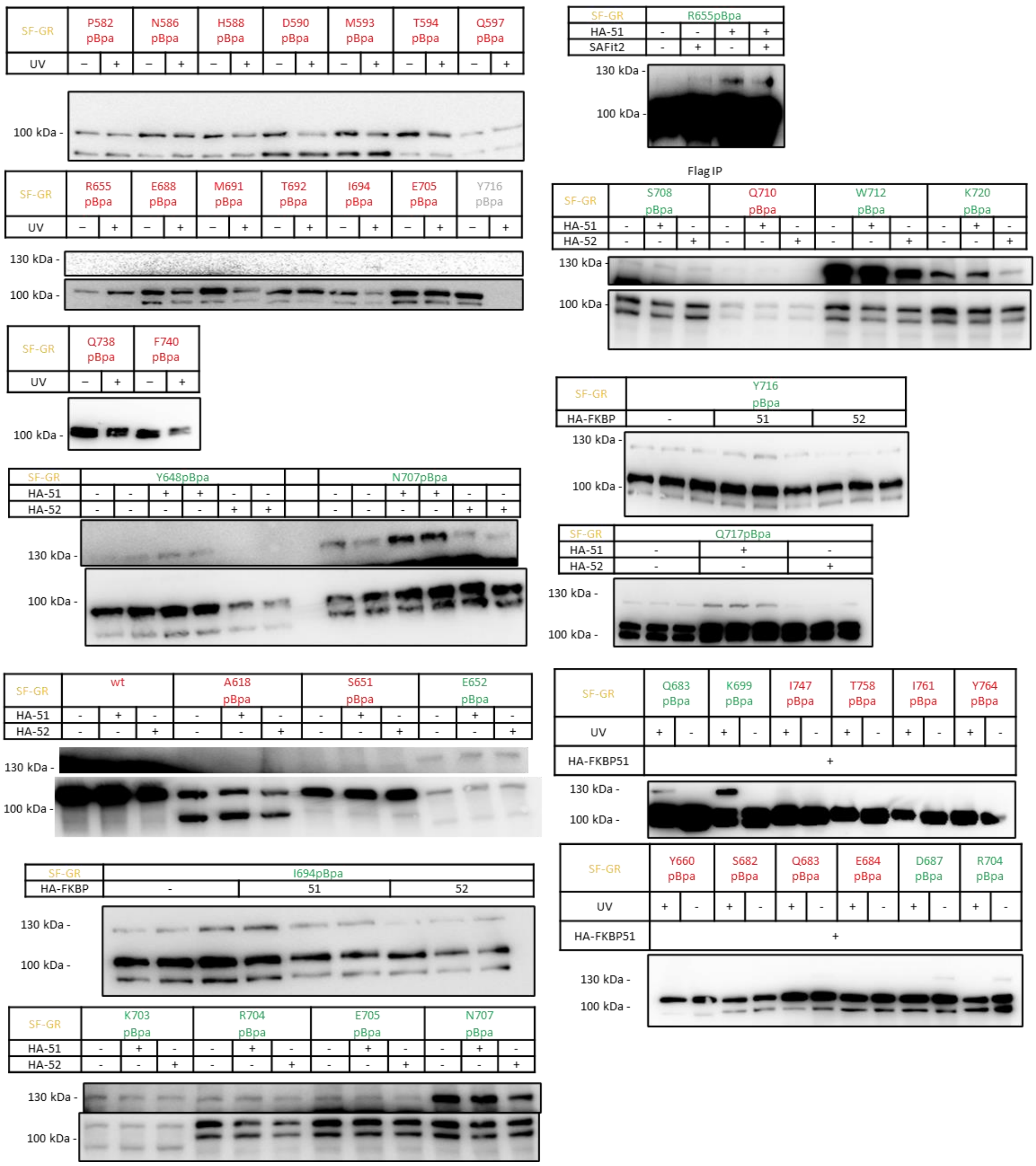


*Supplemental Figure 2:* ***GR-p23 interaction:*** *HEK293 cells co-overexpressing StrepFlag-tagged GR pBpa mutants were UV irradiated and GR→p23 crosslinks were analyzed by Western Blot. Apparent molecular weight (in kDa) is indicated on the left. All blots were stained with an anti-Flag primary antibody.*

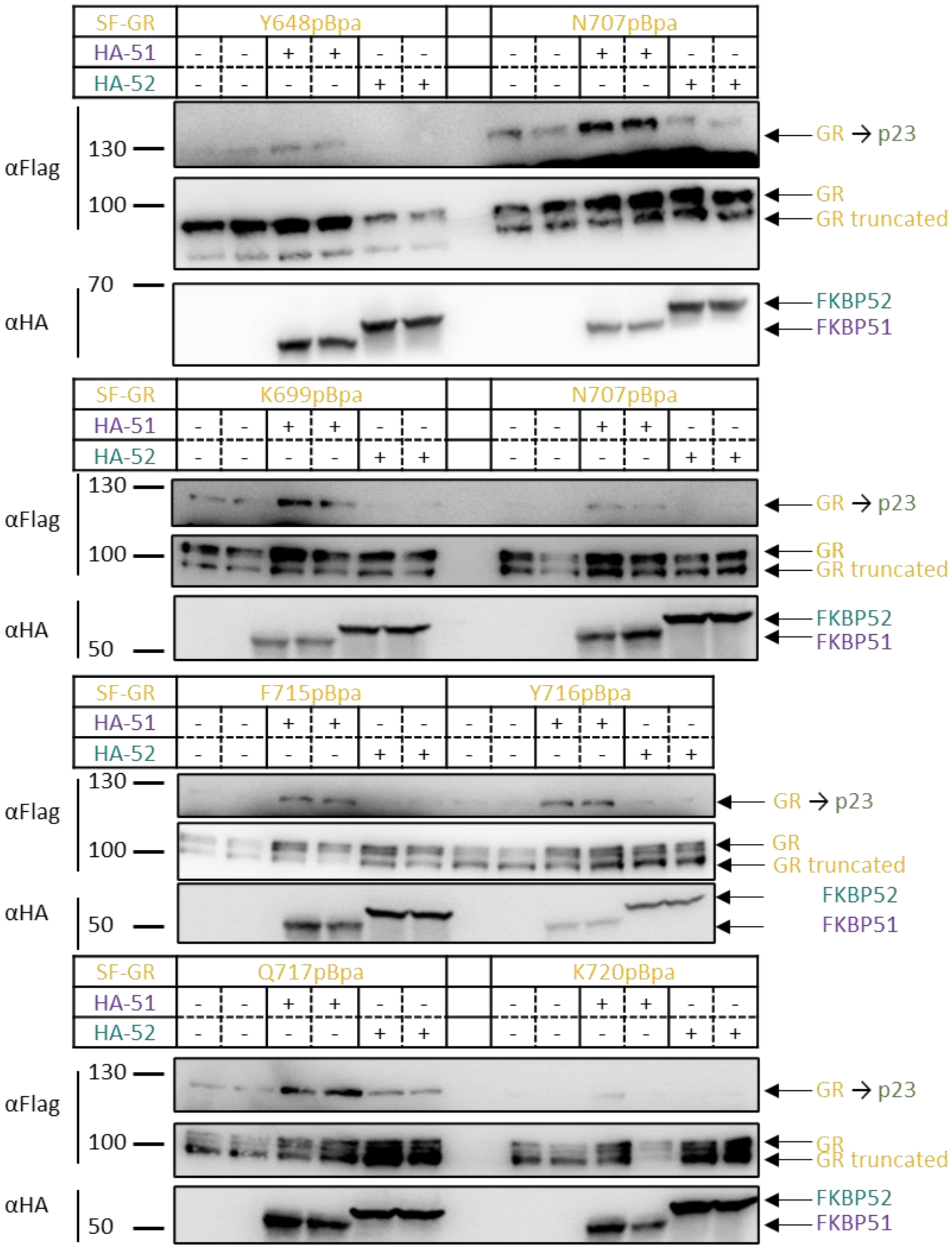


*Supplemental Figure 3:* ***FKBP51 and FKBP52 overexpression influence the GR-p23 interaction:*** *HEK293 cells co-overexpressing Flag-tagged GR pBpa mutants and HA-tagged FKBP51 or FKBP52 were UV irradiated and GR→p23 crosslinks were analyzed by Western Blot in biological duplicates. Apparent molecular weight (in kDa) and primary antibodies are indicated on the left and protein identities are annotated on the right of the blot.*

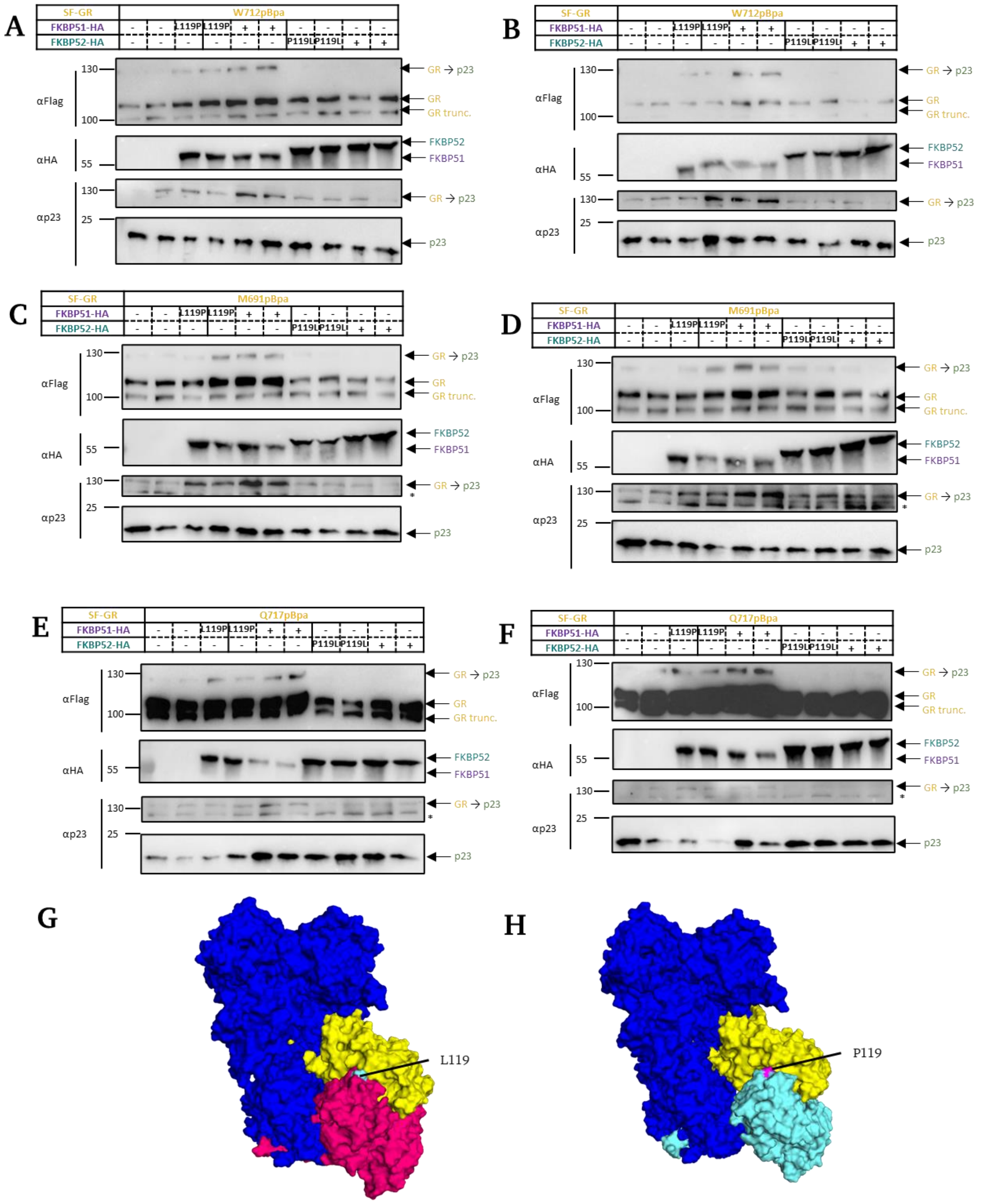


*Supplemental Figure 4:* ***Amino acid swap at position 119 reduces the stabilizing effect of FKBP51 on the GR-p23 interaction in cellulo: A-F:*** *HEK293 cells co-overexpressing Flag-tagged GR pBpa mutants and HA-tagged FKBP51, $FKBP51^{L119P}$, $FKBP52^{P119L}$ or FKBP52 were UV irradiated and GR→p23 crosslinks were analyzed by Western in biological quadruplicates. Protein sizes and primary antibodies are indicated on the left and protein identities are annotated on the right of the blot.* ***G-H****: GR-$Hsp90_2$ complexes with either FKBP51 (**G**, FKBP51 in magenta, L119 marked in cyan, pdb:8FFW) or FKBP52 (**H**, FKBP52 in cyan, P119 marked in magenta, pdb: 8FFV).*

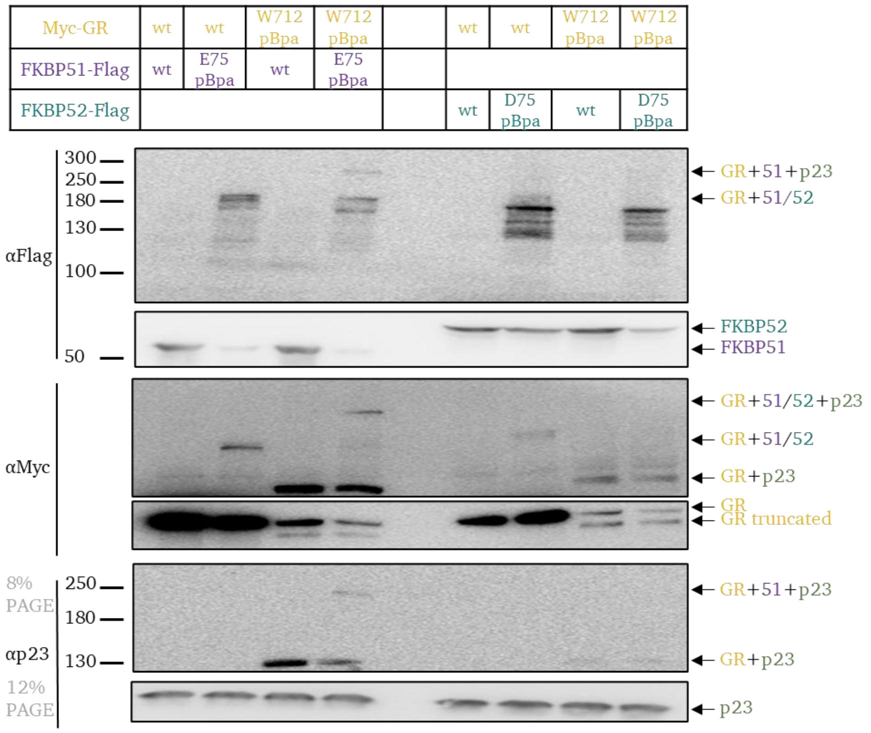


*Supplemental Figure 5:* ***Comparison of the FKBP51-GR-p23 complex with the FKBP52-GR complex****. HEK293 cells co-overexpressing Myc-tagged wildtype GR or GR pBpa mutants, Flag-tagged wildtype FKBP51/FKBP52 or FKBP51/FKBP52 pBpa mutants and HA-tagged p23 were UV irradiated and the photocrosslinks were analyzed by Western Blot. Protein sizes and protein identities are indicated left and right of the blot respectively. Photocrosslinked bands at approximately 300 kDa correspond to the postulated FKBP51→GR→p23 multi-chaperone complex.*

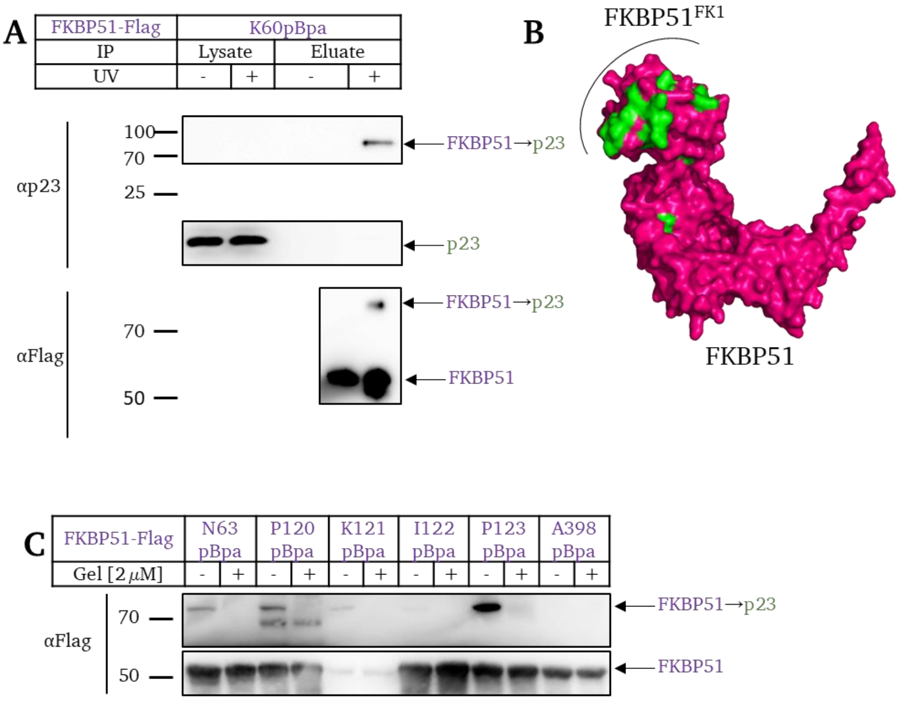


*Supplemental Figure 6: **FKBP51→p23 Crosslinks: A:** Verification of FKBP51→p23 crosslink: HEK293 cells overexpressing Flag-tagged FKBP51 pBpa mutants were UV irradiated and the crosslinks of FKBP51→p23 were purified by Flag IP and subsequently analyzed by Western Blot. Protein sizes and protein identities are indicated left and right of the blot, respectively. **B:** Positions which were tested positive for FKBP51→p23 crosslinks are shown in green on FKBP51 (magenta, 5OMP) **C: FKBP51-p23 interaction is Hsp90 dependent:** HEK293 cells overexpressing Flag-tagged FKBP51 pBpa mutants were treated with 2 μM Geldanamycin for 1 hour, UV irradiated and the crosslinks of FKBP51→p23 analyzed by Western Blot. Protein sizes and protein identities are indicated left and right of the blot, respectively.*
